# Imaging-based temporal dynamics of electrophile-induced NRF2 signaling in liver cells identifies adaptive versus adverse repeated exposure conditions

**DOI:** 10.1101/2025.08.22.671761

**Authors:** Marije Niemeijer, Liesanne Loonstra-Wolters, Bas ter Braak, Hiral Patel, Dawn Yates, Raju P. Sharma, Beate Nicol, Chris Sparham, Sarah Hatherell, Alistair Middleton, Andrew White, Joost B. Beltman, Bob van de Water

**Affiliations:** Division of Cell Systems and Drug Safety, Leiden Academic Centre for Drug Research, Leiden University, Leiden, The Netherlands; Safety, Environmental and Regulatory Science, Unilever, Colworth Science Park, Sharnbrook, Bedford, United Kingdom, MK44 1LQ; Charles River, Cambridge, United Kingdom

**Keywords:** Repeated exposure, oxidative stress response, high content imaging, HepG2 BAC GFP, soft electrophiles

## Abstract

The activation of stress responses upon chemical exposure is critical to restore cellular homeostasis. One of these pivotal pathways is the oxidative stress response activated by NRF2 under control of sensor KEAP1. Soft electrophiles are known to bind with KEAP1 and consequently activate NRF2. However, the understanding of the NRF2 activation dynamics and influence on cellular fate during prolonged repeated exposure conditions is still lacking. Therefore, we accurately mapped the NRF2 activation dynamics upon single or repeated exposures using HepG2 BAC GFP reporters combined with live cell confocal imaging. In a 2D set-up, a concentration-dependent NRF2 activation and subsequent SRXN1 induction upon exposure to sulforaphane or CDDO-Me was observed. The level of NRF2 activation upon sulforaphane exposure highly correlated with cytotoxicity induction. Reactivation of NRF2 upon a second exposure without a recovery period was lower compared to initial exposure, yet resulted in a strong and sustained upregulation of SRXN1. In contrast, inclusion of recovery periods did allow for strong NRF2 and SRXN1 upregulation. In 3D, increased SRXN1 upregulation was observed upon a second exposure with sulforaphane after recovery up to 6 days compared to the first exposure, while similar or lower SRXN1 induction was observed after secondary CDDO-Me treatment. In conclusion, NRF2 activation dynamics and subsequently SRXN1 induction is highly chemical, concentration and exposure duration dependent. Recovery periods allows cells to adapt and initiate a secondary NRF2 response. Together this allowed for further understanding of NRF2 dynamics and cellular fate upon repeated exposures contributing to improving chemical safety testing.

## 1. Introduction

Understanding the regulation of adaptive stress response activation upon (repeated) chemical exposure is critical for improving chemical safety assessment. The activation of specific stress response pathways is frequently an important determinant for adverse outcomes (AOs), which has been recognized within various Adverse Outcome Pathways (AOPs) where early key events often describe such molecular stress (Vinken 2013; Leist et al. 2017). Upon chemical-induced stress, cells will activate these stress response pathways to restore homeostasis and adapt (Weaver et al. 2020). However, when stress conditions are too severe or prolonged due to sustained or repeated exposure, cells can switch towards cell death signaling, and this may lead to adversity (Liu et al. 2013; Hetz and Papa 2018; Hafner et al. 2019). Most *in vitro* experimentation involve single treatment scenarios with short time courses. Yet, human exposure involves repeated dosing for prolonged periods or with different time intervals. Therefore, it is important to accurately map the regulation of these stress responses during repeated exposure scenarios and elucidate the capability to adapt upon multiple insults, an insight that is currently lacking.

Soft electrophiles, present in diet, consumer products or through environmental exposure, are known to induce the oxidative stress response (Yamamoto et al. 2018). Electrophiles are either neutral or positively charged and can react with nucleophiles. Soft electrophiles, such as sulforaphane, can induce the NRF2-mediated oxidative stress response by reacting with cysteine residues of KEAP1 (Hu et al. 2011; Takaya et al. 2012; Yamamoto et al. 2018). This leads to the stabilization and accumulation of NRF2. NRF2 then translocates to the nucleus, where it functions as a transcription factor through binding of antioxidant response elements in the promotor regions of a large battery of NRF2 target genes, such as SRXN1, HMOX1 and NQO1, initiating the antioxidant response to allow for detoxification (Morgenstern et al. 2024). Nevertheless, high levels of exposure to these soft electrophiles may lead to the induction of cell death. Mapping the balance between adaptive and adverse signaling during repeated exposures with soft electrophiles is critical to improve our understanding of cellular adversity. Whether initial activation of the NRF2 response leads to adaptive signaling upon secondary exposures or alternatively sensitizes for cell death induction is still unclear.

To follow the activation of the NRF2 response, we have previously established a BAC GFP reporter panel in hepatocellular carcinoma HepG2 cells using the bacterial artificial chromosome (BAC) recombineering approach (Poser et al. 2008; Wink et al. 2017; Wink et al. 2018). Here, genes of interest were linked to a GFP reporter under control of the endogenous promotor inserted in the genome of the HepG2 cells. For the NRF2 oxidative stress response pathway, reporters were built for KEAP1 as sensor, NRF2 as transcription factor and one of its downstream target genes, SRXN1. The latter target functions as an oxidoreductase and has been identified as a specific and robust biomarker for NRF2 activation in multiple cell types and species (Morgenstern et al. 2024). These HepG2 reporter cells can be grown in 2D conditions, or as 3D spheroids embedded in an extra cellular matrix allowing repeated exposure testing for multiple days in conditions with increased metabolic capacity of cells (Ramaiahgari et al. 2014; Hiemstra et al. 2019). Recently, we evaluated activation of the NRF2 oxidative stress response using these HepG2 reporters upon a first and second exposure to oxidative stress inducers diethyl maleate and tert-butylhydroquinone (Bischoff et al. 2019). In general, the second activation of NRF2 was lower than the first activation, while SRXN1 induction was higher upon a second exposure compared to the first exposure. For other NRF2 inducers sulforaphane, CDDO-Me and nitrofurantoin, a similar further increase in SRXN1 was seen after a second exposure. In this work no recovery period was introduced, which would potentially have allowed cells to adapt to a previous exposure. Moreover, the 2D cell culture conditions prevented repeated exposure for a prolonged period of more than 3 days, and thus did not reflect typical exposure scenarios.

To evaluate the influence of a recovery period between repeated exposures, we made use of these NRF2 response reporters to accurately map the activation dynamics upon exposure to soft electrophiles sulforaphane and CDDO-Me, both potent NRF2 inducers (Thimmulappa et al. 2002; Liby et al. 2005; Hu et al. 2011; ter Braak et al. 2022). The difference in NRF2 activation dynamics upon a second exposure compared to the initial exposure was evaluated using various recovery periods and related to cell death in HepG2 cells cultured in both 2D and 3D, the latter condition allowing for the evaluation of extended exposure periods. Together, our results provide an improved understanding of the activation dynamics of the NRF2 pathway during repeated exposure conditions.

## 2. Materials and Methods

### 2.1 Chemicals and Reagents

Sulforaphane (CAS 142825-10-3) was purchased at Sigma (Zwijndrecht, The Netherlands). CDDO-Me (CAS 218600-53-4) was purchased at Cayman chemical (Ann Arbor, Michigan, USA). Compounds were dissolved in dimethyl sulfoxide (DMSO) from BioSolve (Valkenswaard, The Netherlands) at a 500x stock concentration resulting in a final DMSO concentration of 0.2% (v/v). Aliquots were stored at -20°C.

### 2.2 Cell culture

HepG2 human hepatocellular carcinoma cells were obtained from American Type Culture Collection (ATCC, Wesel, Germany). To monitor the oxidative stress response activation in HepG2 cells upon chemical exposure, NRF2, SRXN1 and HMOX1 were previously tagged with GFP using bacterial artificial chromosome (BAC) recombineering (Poser et al. 2008; Wink et al. 2017; Wink et al. 2018; Hiemstra et al. 2019). HepG2 cells were maintained using Dulbecco’s modified Eagle’s medium (DMEM) supplemented with 10% (v/v) fetal bovine serum (FBS), 25 U/mL penicillin, and 25 µg/mL streptomycin up until passage 20 at 5% CO_2_ and 37°C. Cells cultured in 2D were plated at a density of 10,000 cells per well using Greiner Bio-One black SCREENSTAR 384 well plates (Alphen aan den Rijn, The Netherlands) in DMEM supplemented medium. For 3D spheroid formation, cells were embedded in a layer of 10 µL 5 mg/mL Matrigel matrix basement membrane growth factor reduced (Corning) in Greiner Bio-One black µClear 384 well plates at a density of 1,000 cells per well. Cells were allowed to form spheroids for 24 days in the presence of phenol red free DMEM/F12 high glucose medium supplemented with 10% FBS, 25 U/mL penicillin and 25 µg/mL streptomycin as described earlier (Ramaiahgari et al. 2014; Hiemstra et al. 2019).

### 2.3 Chemical exposures

Cells grown in 2D were exposed at 48 h following plating. Old medium was removed by aspiration, after which 25 µL of medium with propidium iodide (PI) at a final concentration of 100 µM and 25 µL of 2x concentrated compound dilution in supplemented DMEM was added to each well. For repeated exposure scenarios, after 24 or 48 h of exposure medium was replaced with freshly diluted compound.

Cells grown in 3D as spheroids were exposed every day during a 10-day period starting on day 24 of spheroid formation. Every day during exposure old medium was removed by aspiration, after which 25 µL of medium and 25 µL of 2x concentrated compound dilution in supplemented DMEM/F12 was added to each well. PI was added to the medium on days of imaging, namely on day 2, 4, 6, 8 and 10. Specific repeated exposure scenarios were tested with varying length of periods of wash-out, namely 2 days of repeated exposure followed by 2, 4 or 6 days of wash-out and again 2 days of repeated exposure. In addition, continuous repeated exposure for 10 days and 2 days repeated exposure followed by washout of 8 days were also tested. For days during wash-out, old medium was aspirated and replaced with fresh medium.

### 2.4 Confocal imaging

Prior to exposure and imaging, cells were stained with 0.1 µg/mL Hoechst 33342 for 2 h in order to visualize nuclei. To monitor GFP and cell death induction by PI, cells were imaged using a Nikon Eclipse Ti confocal laser microscope (Amsterdam, The Netherlands) with lasers at 408, 488 and 561 nm, automated stage and perfect focus system at 37°C and 5% CO_2_. For HepG2 imaging in 2D or 3D, either the objective Nikon 20x Dry Plan Apo VC NA 0.75, or the objective Nikon 10x Dry Plan Fluor NA 0.3 was used, respectively. Images were taken at a resolution of 512 x 512 pixels. For 2D imaging, every hour images were taken at two locations for each well, where the completion of one acquisition round for all wells within one plate lasted 40 min. For 3D imaging, every two days images were taken acquiring a z-stack with the range of 180 µm with 30 µm steps (i.e., 7 focal planes). One acquisition round for 3D lasted four hours to complete for all wells within one plate.

### 2.5 Quantitative image analyses

Image analysis of HepG2 cultured in 2D was done using CellProfiler version 2.2.0 (Broad Institute, Cambridge, USA) with modules specified previously (Wink et al. 2014). The segmentation of the nuclei was done using an in-house watershed masked clustering algorithm (Di et al. 2012) in ImageJ. Segmentation of cytoplasm was done using the Distance N method with an 8 pixel distance and subtracting the area of the nuclei. GFP intensities were measured by calculating the sum of pixel intensities for each individual cell in the nuclei (NRF2) or cytoplasm (SRXN1). Directly after the second exposure a small increase was observed in GFP intensity in all conditions due to plate handling during the exposure. Therefore, GFP intensities after the second exposure were normalized by subtracting all values with the difference in intensity between the time point just before and immediately after the second exposure. PI positive cells were identified by segmentation of the PI signal and masking the PI objects with the nuclei objects where each PI object should have a minimal overlap of 10% with a nuclei. Output was stored as HDF5 files (Wink et al. 2022).

Image analysis of HepG2 cells cultured as 3D spheroids was done using NIS Elements analysis software (Nikon, Amsterdam, The Netherlands). The maximum pixel intensity across z-stacks for each channel was defined to create a 2D projection. Segmentation of the spheroids was done based on a pixel intensity threshold for the Hoechst signal intensity ranging from 450 to 520 dependent on the signal. The mean GFP pixel intensity was calculated for each spheroid based on all pixels within its segmented 2D projection. Similarly, PI positive cells were identified using a pixel intensity threshold of 1500 and the percentage overlap in the area with identified spheroid objects was determined.

Following both 2D and 3D image analysis, output was further analyzed using in-house R-scripts with the packages ggplot, ggpubr, Rcolorbrewer, plyr, dplyr, data.table and reshape2 in Rstudio (version 1.0.153) (Boston, USA). R-scripts can be retrieved using following Zenodo link.

### 2.6 UPLC-MS/MS for quantification of compound concentrations

To quantify the concentration of compound in the medium and cells during exposure, bioanalytical methods were developed at Charles River Laboratories (Cambridge, UK) based on ultra-performance liquid chromatography-tandem mass spectrometry (UPLC-MS/MS). Following compound exposure in 2D (for 0, 1, 2, 4, 6, 8, 24, 48 and 72 h) and 3D (0, 1, 2, 4, 6, 8 h, 1, 2, 3, 4, 6, 8 and 10 days), 50 µL of medium was collected from each well for the quantification of compound in the medium. To quantify within cells, cells were first washed with 1x PBS and 30 µL of methanol was added. Wells were air-dried at room temperature. When wells were completely dry, 50 µL of distilled water was added and frozen at -80°C prior to shipment to Charles River Laboratories. Samples were prepared by precipitation of matrix proteins with solvents, centrifugation and UPLC-MS/MS analysis of compound recovered in the supernatant.

### 2.7 Statistics

Data is represented as the mean ± SD of three independent biological replicates. For each biological replicate in 2D, the mean was calculated based on imaging of two locations within each well. For 3D, only one location was imaged for every well. Significance was calculated using one-way ANOVA represented as *padj < 0.05, **padj < 0.01, ***padj < 0.001.

## 3. Results

### 3.1 Distinct NRF2 activation dynamics upon exposure to oxidative stress response inducers sulforaphane and CDDO-Me

To evaluate the difference in NRF2 response activation dynamics upon exposure to sulforaphane and CDDO-Me, both potent NRF2 inducers, we exposed hepatocellular carcinoma HepG2 BAC NRF2- and SRXN1-GFP reporter cells, previously developed (Wink et al. 2017; Wink et al. 2018). Using these GFP reporters, signaling dynamics of NRF2 itself as well as one of its downstream target genes, SRXN1, could be followed over time during exposure conditions. Cells were exposed to a broad concentration range of both compounds and imaged every hour for 48 h using confocal microscopy.

As expected, a concentration-dependent induction of nuclear NRF2-GFP was observed upon exposure to sulforaphane in the range of 0.35 to 35 µM (Fig. 1A-B). At all concentrations of sulforaphane, a prominent peak of NRF2-GFP occurred around 2.5 to 10 h upon exposure, after which NRF2-GFP decreased again at later time points. NRF2-GFP was maximally induced at a concentration of 35 µM of sulforaphane (Fig. 1C, middle top panel). In addition, at the concentration of 35 µM, following the initial peak NRF2-GFP levels remained at a heightened level compared to baseline levels and did not decrease anymore at late time points (Fig. 1B). CDDO-Me also exhibited a concentration-dependent induction of NRF2-GFP, however at a lower level than sulforaphane (Fig. 1B). A small peak of NRF2-GFP was observed at early time points, and NRF2-GFP slowly decreased directly after this peak and subsequently remained at the same level at late time points, thereby showing different activation dynamics than sulforaphane. Maximum NRF2-GFP levels were reached at a concentration of 0.46 µM CDDO-Me (Fig. 1C, middle bottom panel). Quantification of the NRF2-GFP response by area-under-the-curve (AUC) and maximum level of NRF2-GFP across time for both sulforaphane and CDDO-Me showed a concentration dependent increase (Fig. 1C, left panels). Interestingly, the maximum NRF2-GFP peak for sulforaphane shifted to later time points with increasing concentrations (Fig. 1C, right top right panel). For CDDO-Me, this fluctuated across different concentrations and remained approximately at the same time point (Fig. 1C, right bottom panel).

**Fig. 1.**
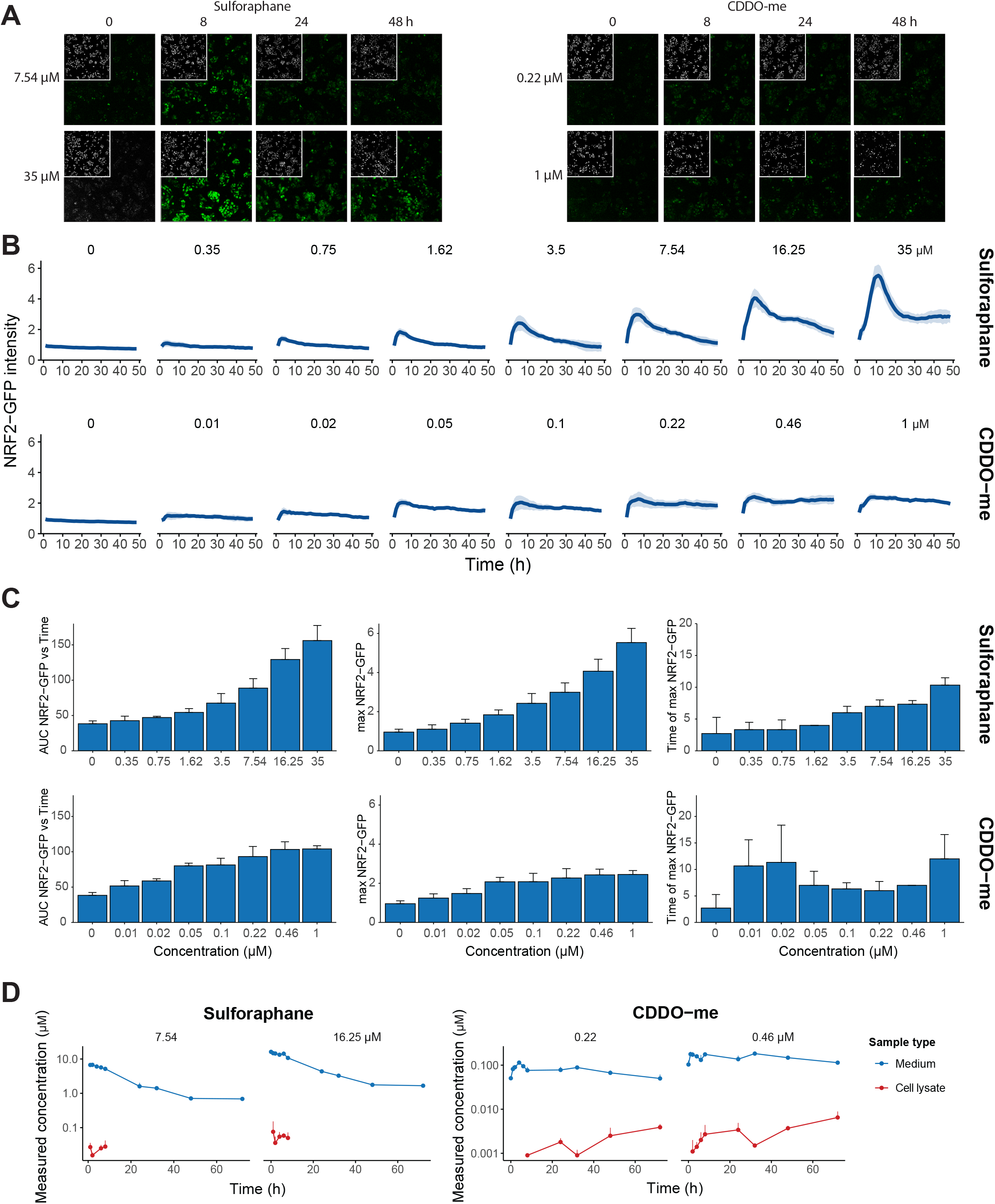
Concentration-dependent activation dynamics of NRF2 upon exposure to sulforaphane and CDDO-Me. HepG2 BAC NRF2-GFP reporter cells were exposed to a broad concentration range of sulforaphane and CDDO-Me and imaged for 48 h live using a confocal microscope with an interval of 1 h. A) Images of HepG2 BAC NRF2-GFP cells stained with Hoechst for nuclei visualization depicted in the top left in grayscale using a Nikon Eclipse Ti microscope and 20x objective. NRF2-GFP signal intensities are depicted in green during exposure at 0, 8, 16, 24, 32, 40 and 48 h time points for three concentrations of sulforaphane and CDDO-Me. B) Activation dynamics of nuclear NRF2-GFP during exposure with sulforaphane and CDDO-Me. C) Quantifications of the area under the curve (AUC), maximum NRF2-GFP intensity over time and the exposure time at the maximum of NRF2-GFP upon exposure with sulforaphane and CDDO-Me. D) The measured concentration of sulforaphane and CDDO-Me using UPLC-MS/MS in the medium and within cell lysates during exposures. Data presented as the mean of N=3, error bars represented as the SD.

Given the differences in NRF2 activation dynamics between sulforaphane and CDDO-Me, next, we used UPLC-MS/MS to measure the concentration of both compounds intracellularly and in the medium during exposures at two concentrations (Fig. 1D). Interestingly, sulforaphane levels markedly dropped both in the medium and in the cells during 72 h, in the latter case to levels lower than the detection limit. In contrast, detected CDDO-Me slightly increased at early time points and remained at a similar level at late time points. In conclusion, compound kinetics and NRF2 activation dynamics exhibit a compound-dependent pattern over time, suggesting that the difference in kinetics of these compounds could partially be responsible for the difference in NRF2 activation dynamics over time, where CDDO-Me remains at a stable level while sulforaphane is less stable.

### 3.2 Downstream NRF2 target gene SRXN1 activation dynamics

Activation dynamics of SRXN1, a downstream target gene of NRF2, was evaluated upon exposure for 48 h to sulforaphane and CDDO-Me in HepG2 BAC SRXN1-GFP reporter cells (Fig. 2). A concentration-dependent induction occurred upon exposure to both compounds (Fig. 2A-B). For sulforaphane, a maximum SRXN1-GFP level was reached at a concentration of 16.25 µM. At 35 µM of sulforaphane, the SRXN1-GFP level was lower than at 16.25 µM, suggesting that the threshold of cellular toxicity had been reached. The time point at which SRXN1-GFP levels were maximal were approximately 30 hours of exposure, and SRXN1-GFP decreased again at later timepoints. For CDDO-Me, the highest level of SRXN1-GFP was reached at 0.46 µM, whereas it was lower at a concentration of 1 µM. During 48 hours of continuous exposure to CDDO-Me, SRXN1-GFP kept increasing in contrast to sulforaphane. This is in concordance with the distinct NRF2 activation dynamics and kinetic differences between sulforaphane and CDDO-Me (Fig. 1). Indeed, peak levels of NRF2- and SRXN1-GFP correlated well for sulforaphane (Pearson correlation of 0.73) (Fig. 2C). Only at the highest concentration of sulforaphane, peak SRXN1-GFP levels decreased compared to the second-highest concentration, while NRF2 levels remained high. Also for CDDO-Me, NRF2- and SRXN1-GFP correlated well, although the Pearson correlation was slightly less strong (0.63). At high concentrations, peak levels of NRF2 remained similar while SRXN1 levels increased across the investigated concentration range. In conclusion, the differential kinetics and NRF2 activation dynamics between sulforaphane and CDDO-Me resulted also in differential dynamics of downstream target SRXN1.

**Fig. 2.**
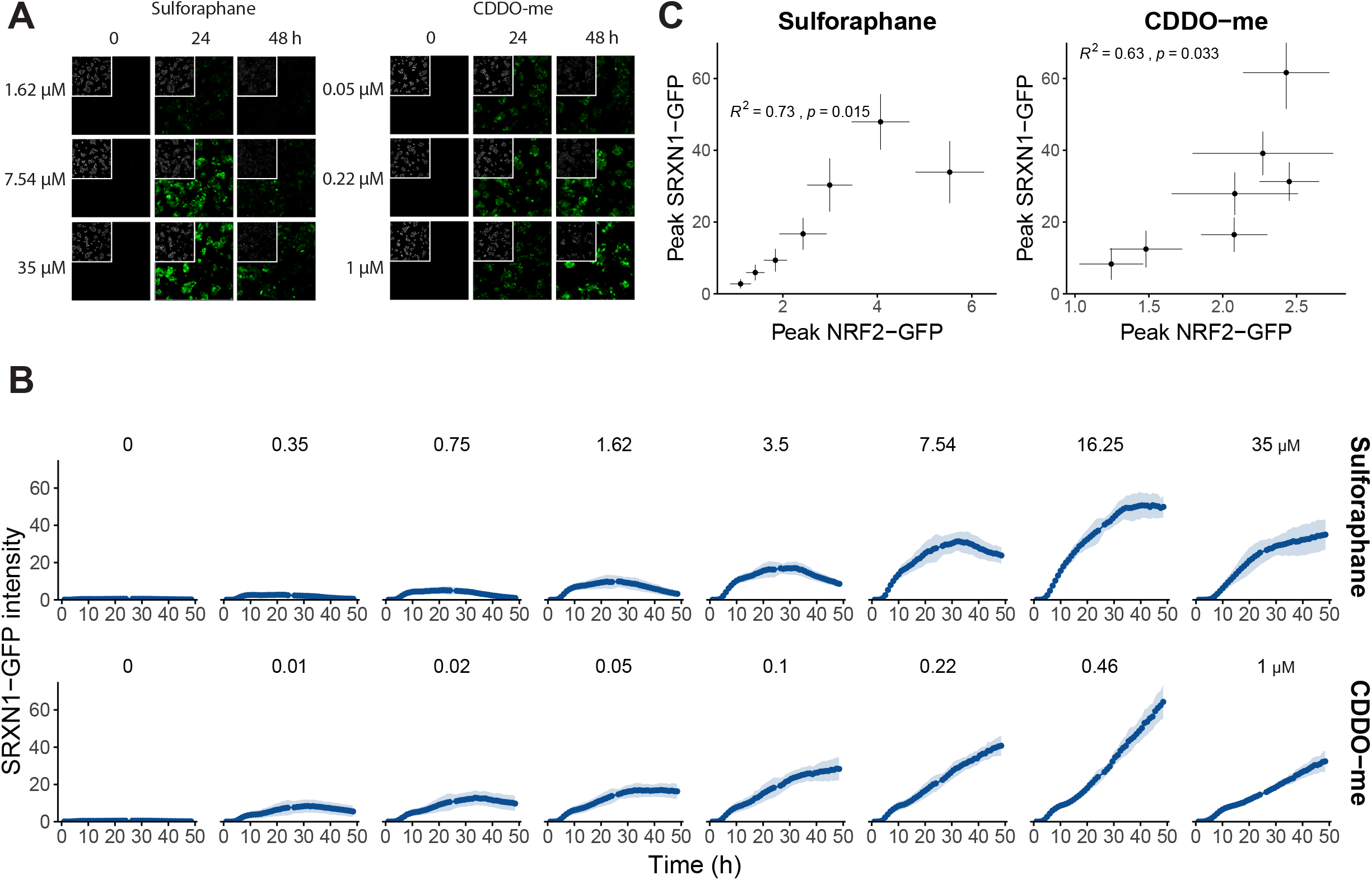
Activation dynamics of NRF2 target gene SRXN1 upon exposure to sulforaphane and CDDO-Me. HepG2 BAC SRXN1-GFP reporter cells were exposed to a broad concentration range of sulforaphane and CDDO-Me and imaged for 48 h live using a confocal microscope with an interval of 1 h. A) Confocal images of HepG2 BAC SRXN1-GFP cells stained with Hoechst for nuclei visualization upon exposure to sulforaphane and CDDO-Me for 24 or 48 h at three different concentration levels. SRXN1-GFP signal intensities are depicted in green. B) Activation dynamics of cytoplasmic SRXN1-GFP upon exposure to sulforaphane and CDDO-Me for 48 h. C) Correlation between NRF2- and SRXN1-GFP activation upon sulforaphane (left panel) and CDDO-Me (right panel) exposure. The square of the Pearson correlation (R^2^) and p-value are depicted. Data presented as the mean of N=3, error bars represented as the SD.

### 3.3 Correlation of NRF2 activation with cell death induction

To evaluate the correlation between NRF2 response activation dynamics and cellular injury, we quantified the number of cells during exposure (Fig. 3A, S1). Here, a concentration-dependent decrease in cell number growth during exposure was evident for both sulforaphane and CDDO-Me. At the highest concentrations, the total cell number remained the same or became lower over time for sulforaphane and CDDO-Me, respectively, indicating cell death induction or a decrease in proliferation. This drop in cell number correlated strongly with the cumulative NRF2-GFP induction for sulforaphane, but this correlation was less pronounced for CDDO-Me (Fig. 3B). Similarly, the cumulative NRF2-GFP induction correlated more strongly with the fraction of propidium iodide (PI) positive cells (a dye indicating cell death) for sulforaphane than for CDDO-Me (Fig. 3C, S2). Correlations between SRXN1-GFP and total or PI-positive cell numbers were lower than for NRF2-GFP (Fig. 3D, E, S2). These analyses, for sulforaphane in particular, suggest that NRF2 is a potential determinant of cell death at increasing concentrations.. For sulforaphane, cell death gradually increased with applied compound concentration, whereas for CDDO-Me a sharper concentration threshold existed above which cells started to die.

**Fig. 3.**
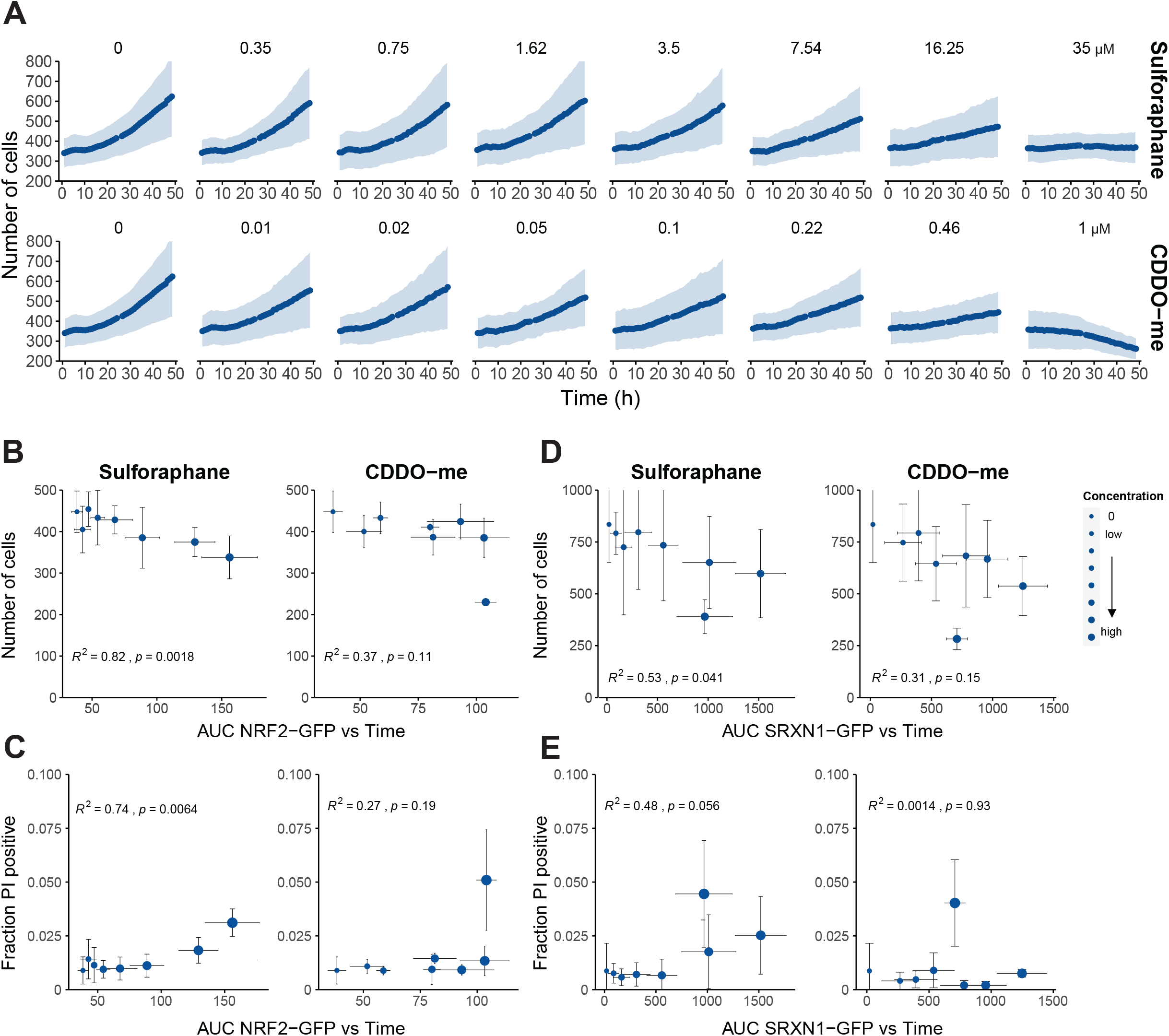
Correlation between NRF2 response activation and cell death induction upon exposure to sulforaphane and CDDO-Me. HepG2 NRF2- and SRXN1-GFP reporter cells were exposed to a broad concentration range of sulforaphane and CDDO-Me. Cells were live imaged for 48 h using confocal microscopy with a time interval of 1 h. Total number of cells and cell death was evaluated by quantifying the mean fraction of propidium iodide (PI) positive cells of both HepG2 NRF2- and SRXN1-GFP reporter cells (each N=3). A) Total number of cells during exposure until 48 h with sulforaphane and CDDO-Me. B-E) Correlation between area under the curve (AUC) across time of NRF2-GFP (B, C) or SRXN1-GFP (D, E) and number of cells (B, D) or fraction of PI positive cells (C, E) upon exposure to sulforaphane and CDDO-Me (concentrations depicted as symbol size). The square of the Pearson correlation (R^2^) and p-values are depicted within each graph. Data presented as the mean of N=3, error bars and shading represented as the SD.

### 3.4 Secondary NRF2 response activation during repeated exposures

We previously found that the NRF2 response is differentially regulated upon a secondary exposure when already activated by an initial exposure and without a recovery period using diethyl maleate and CDDO-Me as inducers (Bischoff et al. 2019). As a next step, we exposed HepG2 BAC NRF2- and SRXN1-GFP reporter cells to sulforaphane and CDDO-Me in a broad concentration range for 24 h followed by a second exposure for 24 h (Fig. 4). We captured GFP intensities every hour with confocal microscopy from 0 to 48 h and compared the NRF2 and SRXN1 activation dynamics upon the first exposure with the dynamics upon a second exposure.

**Fig. 4.**
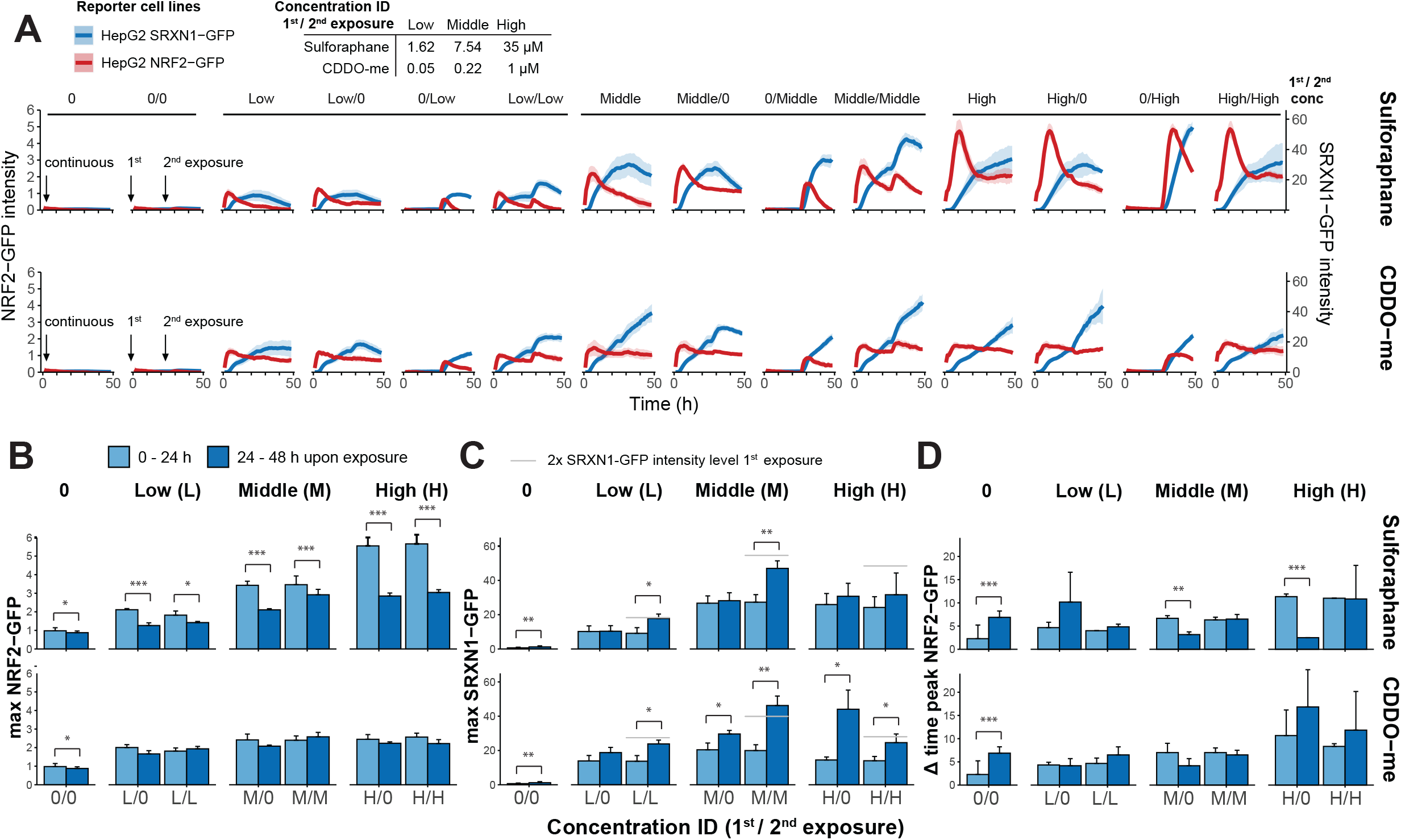
Influence of pre-existent NRF2 activation on secondary NRF2 activation dynamics during repeated exposure scenarios. HepG2 BAC NRF2- and SRXN1-GFP cells were exposed at different concentration levels (0, low, middle and high) to sulforaphane (0, 1.62, 7.54 or 35 µM, respectively) or CDDO-Me (0, 0.05, 0.22 or 1 µM, respectively) followed by a second exposure at the 24 h time point. Cells were imaged live using a confocal microscope with a time interval of 1 h for 0 to 48 h during exposures. A) Quantification of the NRF2- (red) and SRXN1-GFP (blue) intensity during first (0 – 24 h) and second (24 – 48 h) exposures with different combinations of concentration levels (0, low, middle or high) of sulforaphane and CDDO-Me. B-C) Maximum NRF2-GFP (B) and SRXN1-GFP (C) intensity after first (0 – 24 h, light blue bars) and second (24 – 48 h, dark blue bars) exposure at different concentration levels of sulforaphane and CDDO-Me (0, low, middle or high). Grey lines in (C) represent the level of two fold of the maximal SRXN1-GFP intensity during first exposure (0 – 24 h) as reference. D) Length of time interval of NRF2-GFP peak after first (0 – 24 h) and second (24 – 48 h) exposure at different concentration levels of sulforaphane and CDDO-Me (0, low, middle or high). Data presented as the mean of N=3, error bars and shading represented as the SD. Significance was calculated using one-way ANOVA represented as *padj < 0.05, **padj < 0.01, ***padj < 0.001.

For sulforaphane, the second induction of NRF2 was lower compared to the initial induction at the low and middle concentrations, yet a clear second activation peak of NRF2 could be observed (Fig. 4A, top panels; Fig. 4B). For the highest concentration of sulforaphane, NRF2 was minimally induced upon the second exposure and remained at the same level. A similar pattern could be observed for SRXN1-GFP upon sulforaphane exposure: At low and middle concentrations SRXN1 was much more strongly induced upon a second exposure while at high concentration minimal further induction occurred. A second exposure with CDDO-Me resulted in similar levels of NRF2-GFP compared to the first exposure (Fig. 4A, bottom panels; Fig. 4C).

Despite similar levels of NRF2-GFP, a second exposure with CDDO-Me resulted in substantially higher SRXN1-GFP levels compared to conditions without second exposure (continuous) or upon the first exposure. The timing of the peak of NRF2 remained similar when comparing activation upon a first or second exposure for both compounds at all concentrations (Fig. 4D). In conclusion, consistent with our earlier findings for repeated exposures to DEM (Bischoff et al. 2019), second exposure to either sulforaphane or CDDO-Me led to further SRXN1 induction, whereas NRF2 dynamics exhibited a similar or lower induction as during the first exposure.

### 3.5 Effect of recovery period on oxidative stress response re-activation

To evaluate if the oxidative stress response is similarly activated upon a second exposure when cells are allowed to recover for a certain amount of time, we exposed HepG2 BAC NRF2- and SRXN1-GFP cells for 24 h with either sulforaphane or CDDO-Me followed by a 24 h recovery period with fresh medium and a second exposure with the same compound for 24 h. Cells were imaged every hour for 72 h to follow the difference in NRF2 activation dynamics (Fig. 5).

**Fig. 5.**
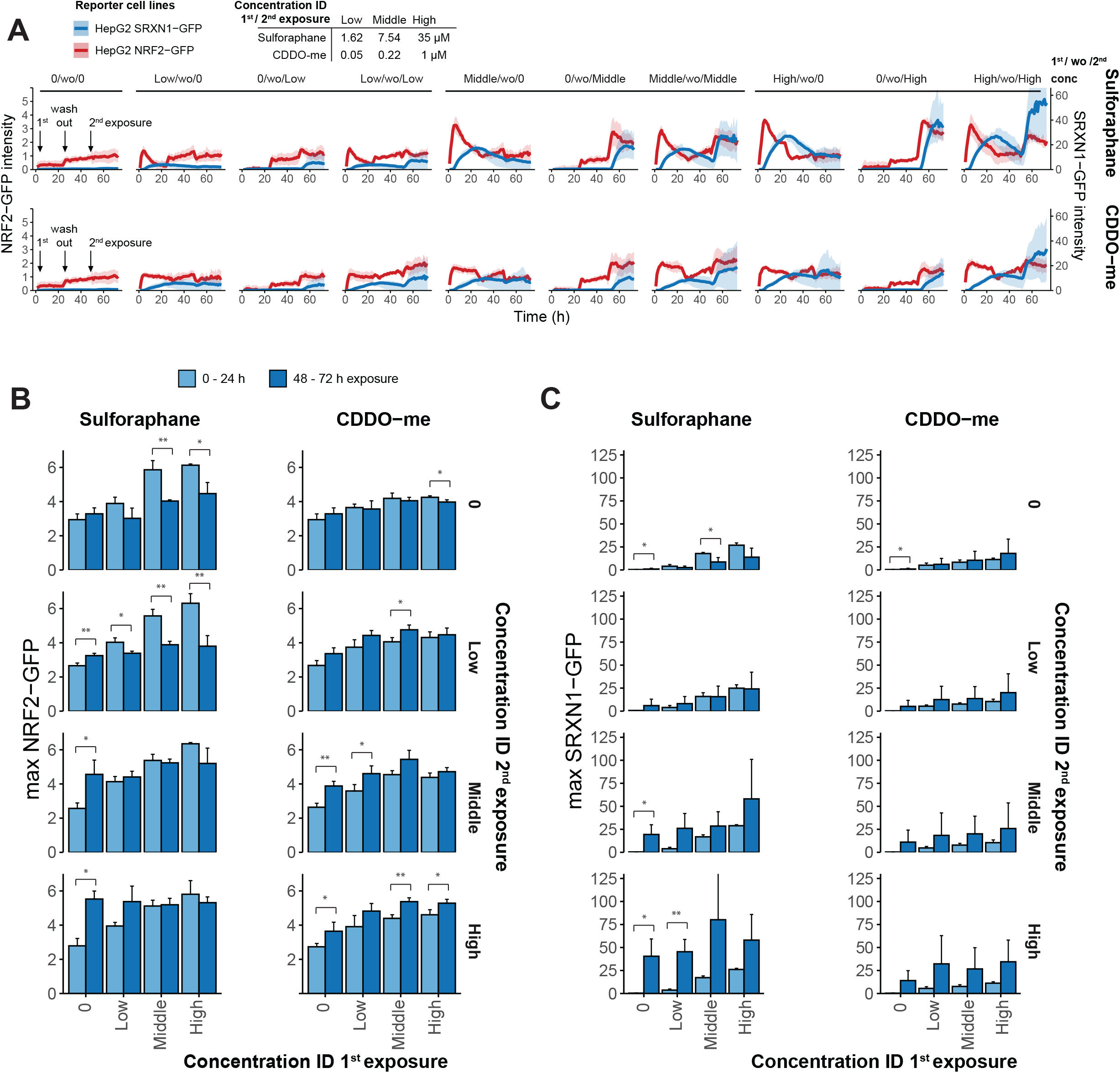
Effect of recovery period on NRF2 response re-activation during repeated exposures. HepG2 BAC NRF2- and SRXN1-GFP cells were exposed for 24 h at different concentration levels (0, low, middle, high) to sulforaphane (0, 1.62, 7.54 or 35 µM, respectively) and CDDO-Me (0, 0.05, 0.22 or 1 µM, respectively)followed by a recovery period for 24 h and a second exposure at the 48 h time point for 24 h. Cells were imaged live using a confocal microscope with a time interval of 1 h for 0 to 72 h during exposures. A) Quantification of the NRF2- (red) and SRXN1-GFP (blue) intensity during first exposure (0 – 24 h), recovery period (24 – 48 h) and second exposure (48 – 72 h) with different combinations of concentration levels of sulforaphane and CDDO-Me (0, low, middle or high). B-C) Maximum NRF2-GFP (B) and SRXN1-GFP (C) intensities after first (0 – 24 h, light blue bars) and second exposure (48 – 72 h, dark blue bars) at different concentration levels of sulforaphane and CDDO-Me (0, low, middle or high). Data presented as the mean of N=3, error bars represented as the SD. Significance between first and second exposures was calculated using one-way ANOVA represented as *padj < 0.05, **padj < 0.01, ***padj < 0.001.

NRF2-GFP was re-activated upon a second exposure with sulforaphane after a recovery period, albeit with a lower peak compared to the first exposure (Fig. 5A, top, red lines, and 5B, left). This was also observed when no recovery period was included (Fig. 4). However, now a clear second peak was observed also at the highest concentration of sulforaphane which was not the case in the absence of a recovery period. Also, SRXN1-GFP was clearly induced upon a second exposure at all three concentration levels of sulforaphane, leading to higher intensity levels at the second exposure compared to the first exposure for the middle and high concentrations (Fig. 5A, top, blue lines and 5C, left). In general, GFP intensities were more variable upon a second exposure after recovery than in conditions without recovery.

Upon a second exposure with CDDO-Me following a recovery period, NRF2-GFP was further induced compared to the initial activation, although a clear secondary peak did not occur (Fig. 5A, bottom, red lines and 5B, right). SRXN1-GFP was also further induced upon a secondary exposure following a recovery period (Fig. 5A, bottom, blue lines and 5C, right), although this did not reach statistical significance due to large variability across replicates. Interestingly, without a recovery period, the highest concentration did not lead to reactivation of NRF2-GFP and SRXN1-GFP for both compounds, while the presence of a 24 h recovery period in between did result in a clear reactivation upon a second exposure. This suggests that 24 h is sufficient recovery time for cells to adapt. Alternatively, cells that died during first exposure might have been washed away at the start of the recovery phase when medium was replaced.

### 3.6 NRF2 response activation during repeated exposure for 10 days using HepG2 spheroids

To study the influence of prolonged recovery periods (multiple days) on the NRF2 response activation, we grew HepG2 BAC SRXN1-GFP reporter cells as 3D spheroids in a Matrigel layer, allowing for repeated and prolonged exposures up to 10 days (Hiemstra et al. 2019). Following spheroid formation, spheroids were exposed for 10 days, with a repeated exposure every day, or 2 days repeated with varying recovery periods (2, 4 or 6 days) followed by a second 2-day repeated exposure in a broad concentration range of sulforaphane and CDDO-Me (Fig. 6A-B). Cell death induction was only observed with 10 day repeated exposure and highest concentration of CDDO-Me (Fig. S3).

**Fig. 6.**
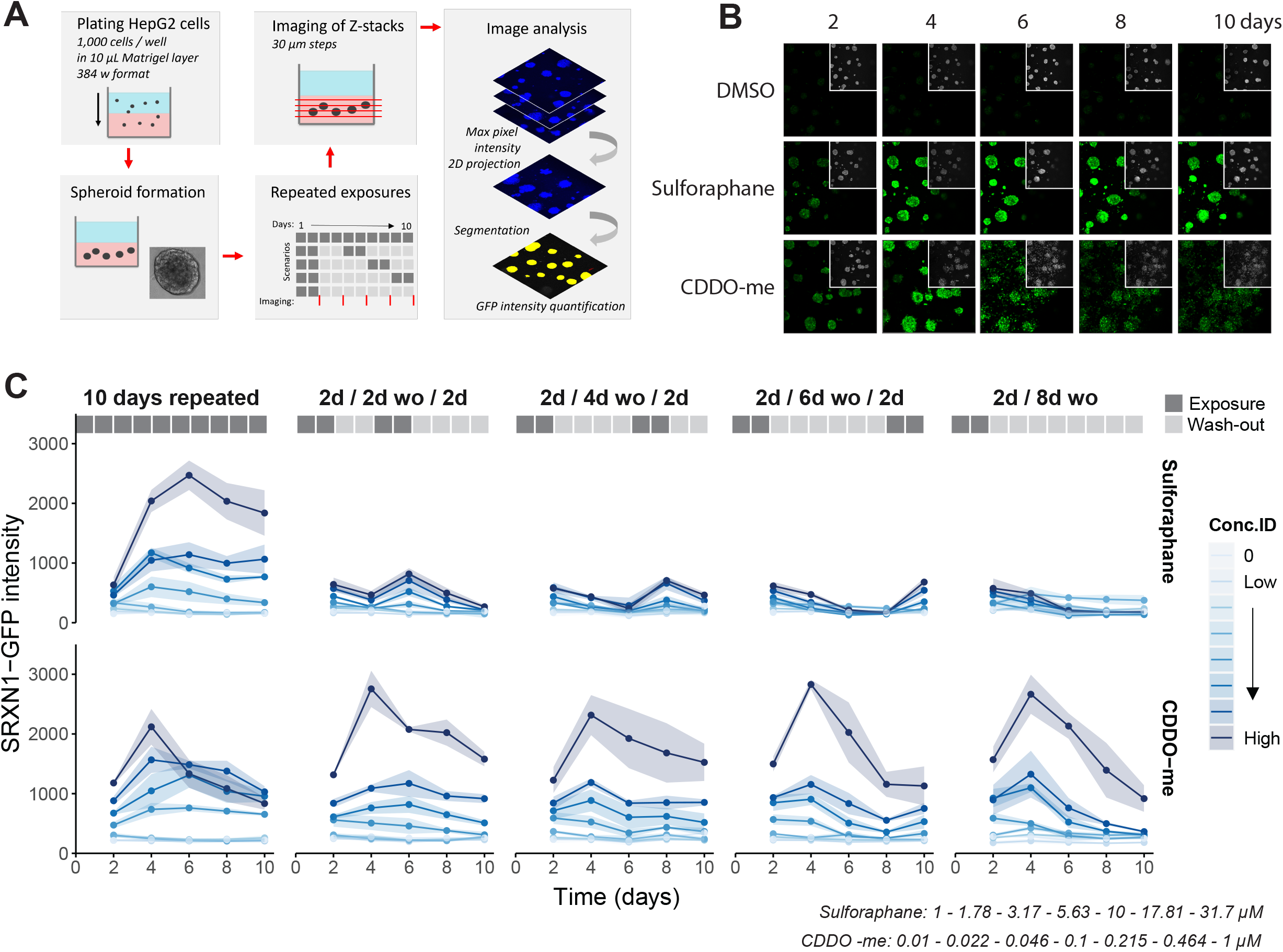
SRNX1 upregulation upon repeated exposures for 10 days in HepG2 3D spheroids. HepG2 BAC SRXN1-GFP cells were allowed to form spheroids for 21 days in a layer of Matrigel, and subsequently exposed every day for 10 days with either sulforaphane or CDDO-Me (using different exposure scenarios). Every two days, spheroids were imaged using a confocal microscope with a 10x objective, creating a z-stack with 30 µm steps. A) Schematic overview of experimental setup and image analysis strategy. B) Images of HepG2 SRXN1-GFP spheroids at 2, 4, 6, 8, and 10 days of exposure to either sulforaphane or CDDO-Me. Nuclei were visualized using Hoechst staining and depicted as grayscale images at top right corners. C) Quantification of SRXN1-GFP during repeated exposure scenarios for sulforaphane and CDDO-Me at 2, 4, 6, 8 and 10 days of exposure. Spheroids were exposed either every day for 10 days, or for 2 days of repeated exposure followed by 2, 4 or 6 days wash-out and 2 days of repeated exposure. Data presented as the mean of N=3, error bars represented as the SD.

Upon 10-day repeated exposure with sulforaphane, SRXN1-GFP was induced in a concentration-dependent manner (Fig. 6C, leftwards panels). From 4 days onwards, SRXN1-GFP levels remained approximately at the same level as for earlier time points, except for the highest concentrations of sulforaphane for which SRXN1-GFP clearly decreased at late stages. A similar pattern occurred for CDDO-Me, i.e. SRXN1-GFP induction reached a maximal level after 4 days that was approximately maintained only for low concentrations.

When a recovery period was applied, SRXN1-GFP levels were slightly higher, although not significant (ρ_adj_ = 0.1297), after the second period of 2-day repeated exposure to sulforaphane compared to those following the initial 2-days repeated exposure period with sulforaphane (Fig. 6C, top). This was the case for all recovery period durations and was in concordance with the results in 2D with a recovery period of 24 h (Fig. 4). For low concentrations of CDDO-Me, a recovery period of up to 6 days led to a continued increase in SRXN1-GFP during the (initial phase of the) recovery period, whereas upon a second 2-day repeated exposure period the levels of SRXN1-GFP remained approximately the same. For the highest concentration of CDDO-Me, SRXN1-GFP even had the tendency to decrease following the second repeated exposure. Overall, for sulforaphane the SRXN1-GFP induction thus followed a similar pattern upon repeated exposure in 3D HepG2 spheroids as in 2D settings, with repeated exposure leading to additional SRXN1-GFP induction even after a recovery period. For CDDO-Me, the SRXN1-GFP induction appeared less sensitive to secondary triggering in 3D compared to 2D.

## 4. Discussion

In this study, we determined the difference in NRF2 response activation upon repeated exposures to soft electrophiles sulforaphane and CDDO-Me. To study the activation dynamics, we combined high-content confocal imaging with HepG2 BAC GFP reporters for the NRF2 pathway in a 2D setting and in a 3D spheroid setup to allow for long-term repeated exposure scenarios. Concentration-dependent activation of NRF2 and SRXN1 occurred upon exposure to sulforaphane and CDDO-Me, both with distinct activation patterns that are likely partially due to differences in kinetics. In related work, we further investigated the relation between NRF2 activation and SRXN1 induction based on these novel experimental data using mathematical modeling (Sharma et al. 2025). In this way, we showed that besides the compound-dependent kinetics that drive NRF2 activation, also the relation between NRF2 and SRXN1 is compound dependent.

In the current study, the level of sulforaphane-mediated NRF2 induction correlated strongly with cell death induction. Secondary exposures led to minor additional NRF2 induction, but SRXN1 was strongly further induced at non-cytotoxic concentrations for both chemicals. In contrast, recovery periods enabled also a strong secondary NRF2 reactivation upon second exposure, on top of the strong SRXN1 activation that occurred both with and without recovery. Finally, using 3D HepG2 spheroids, a second repeated exposure with sulforaphane led to slightly enhanced SRXN1 induction while with CDDO-Me second exposure the SRXN1 levels typically stayed at similar levels or decreased. Together, these results highlight the capability of reactivating the NRF2 response upon secondary exposures following a period of recovery.

The soft electrophiles sulforaphane and CDDO-Me led to distinct oxidative stress activation dynamics, where sulforaphane strongly induced the NRF2 response initially, followed by a rapid decrease. In contrast, CDDO-Me resulted in prolonged NRF2 response activation also at late time points. Possibly this is related to differences in kinetics, since we showed that sulforaphane was less stable than CDDO-Me, i.e., the sulforaphane concentration decreased rapidly in both medium and cells. This is consistent with prior measurements, estimating the half-life of sulforaphane as 2.2 h and for CDDO-Me as approximately 39 h (Hanlon et al. 2008; Hong et al. 2012). Similar differences in NRF2 activation dynamics between sulforaphane and CDDO-Me were reported by ter Braak et al. (ter Braak et al. 2022).

Activation levels of NRF2 were correlated with cell death induction upon specifically sulforaphane exposure. Multiple studies have shown the cytoprotective capabilities of sulforaphane towards oxidative stress by inducing the adaptive NRF2 pathway (Kubo et al. 2017; Wei et al. 2021). However, at high concentrations of sulforaphane other secondary stress response pathways, such as ER stress signaling, may be activated contributing to the switch from adaptive signaling to cytotoxicity (ter Braak et al. 2022; Wijaya et al. 2024). Copple et al. (2019) showed that NRF2 activation, based on TG-GATES transcriptomics data, was highly indicative for intrinsic biochemical reactivity and chemical stress. However, using NRF2 activation alone led to a low sensitivity for drug-induced liver injury (DILI) identification, suggesting that the combination of multiple stress response related factors may contribute to DILI development. Indeed, Souza et al. (2017) found that liver injury-inducing compounds affected multiple stress pathways, such as NRF2, ER stress and DNA damage signaling, while non-liver injury inducing compounds specifically only affected inflammatory NF-kB signaling. Another study by Liu et al. (2022) based on TG-GATES transcriptomics data showed significant enrichment of the expression and regulon activity of NRF2 but also of ATF4, a transcription factor related to ER stress signaling, and of transcription factors related to other stress signaling preceding adverse histopathological changes. Their work provided strong evidence for the time-dependent role of multiple stress pathways in drug-induced liver injury development. Therefore, the balance between adaptivity and the development of adversity upon chemical-induced NRF2 activation is likely highly dependent on the chemical exposure level, duration and number of insults, because these factors jointly determine whether secondary cellular stresses are induced as well.

Allowing cells to recover after chemical insult potentially enables reactivation of the adaptive stress response pathway, thereby preventing the attainment of a breaking point leading to adversity. Indeed, we demonstrated that the NRF2 response was reactivated in a similar manner as during the primary exposure following a recovery period. Without a recovery period there was less reactivation. Furthermore, even though NRF2 was minimally reactivated in conditions without recovery, SRXN1 was highly increased upon a second exposure. This was especially the case for CDDO-Me in our 2D set-up, where NRF2 levels were lower than for sulforaphane but still led to high SRXN1 levels. Potentially, the high stability of CDDO-Me leads to a longer period of NRF2 levels that are sufficiently high to promote further SRXN1 transcription. This was also observed in previous work by Bischoff et al. (2019) where CDDO-Me and diethyl maleate led to less NRF2 reactivation after a second exposure while SRXN1 was still further induced (Bischoff et al. 2019). A study by Mathew et al. (2014) showed similar accumulation of HMOX1 and NQO1, both NRF2 target genes, in fibroblasts upon repeated exposure for three days with sulforaphane.

HepG2 3D spheroids models allowed us to investigate the adaptive response across a prolonged period of ten days, which is more relevant for *in vivo* repeated dosing scenarios. Moreover, unlike HepG2 cells grown in a 2D setting, HepG2 cells within spheroids are quiescent, non-proliferative and express biomarkers that are associated with a more mature hepatocyte phenotype in 3D Matrigel culture (Ramaiahgari et al. 2014; Hiemstra et al. 2019). Repeated dosing for ten days led to a sustained NRF2 activation as apparent from the high levels of SRXN1-GFP expression. Yet, washout of sulforaphane led to baseline levels of SRXN1-GFP after 6 days of recovery with normal SRXN1-GFP induction on day 8. Washout followed by secondary exposure to CDDO-Me has less of an effect on SRXN1-GFP than sulforaphane, with high concentrations of CDDO-Me during secondary exposure even leading to a decreasing expression of SRXN1-GFP. These data indicate the importance of understanding temporal dynamics of NRF2 activity induced by xenobiotics and the relevance of washout and repeated exposure treatment for prolonged time periods. Our results illustrate that different compounds may have different long-term temporal dynamics which may be favorable for sustained pharmacological efficacy but at the same time may increase risks for adverse responses. Although SRXN1 was identified as a robust and specific marker for NRF2 activation (Morgenstern et al. 2024), other critical NRF2 downstream target genes, such as NQO1 or TXNRD1, may show different signaling dynamics upon repeated exposure scenarios affecting cellular fate. Therefore, future studies investigating the role of NRF2 reactivation and possible other regulators of the accumulation of a panel of downstream target genes is necessary to improve understanding of its tight regulation and the influence on the balance between adaptivity and development of adversity during repeated exposure scenarios.

Taken together, regulation of the NRF2 response activation during repeated exposure scenarios is highly dependent on the chemical, exposure duration, xenobiotic concentration and recovery periods applied. Recovery periods allow cells to adapt and circumvent adverse signaling upon a second exposure. Knowledge on the consequences of repeated exposures on adaptive signaling and cellular outcome hopefully will aid in improving chemical safety testing strategies.

## 5. Funding

This work was supported by a Unilever SEAC research grant and the European Commission Horizon 2020 program projects EU-ToxRisk (grant agreement number 681002) and the RISK-HUNT3R (grant agreement number 964537).

## 6. Data availability

Quantification of imaging data can be found using following link. R-scripts are available using following Zenodo link.

## 7. Conflict of interest

The HepG2 BAC GFP reporters are currently licensed to Toxys. BtB is currently employed at this company.

## 8. CRediT authorship contribution statement

**MN:** Writing – original draft, Visualization, Methodology, Investigation, Formal analysis. **LLW:** Writing - Review & Editing, Methodology, Investigation, Formal analysis. **BtB:** Writing - Review & Editing, Methodology, Investigation, Formal analysis. **HP:** Writing - Review & Editing, Methodology, Investigation, Formal analysis. **DY:** Writing - Review & Editing, Methodology, Resources. **RPS:** Writing - Review & Editing, Methodology, Formal analysis. **BN:** Writing - Review & Editing, Conceptualization. **CS:** Writing - Review & Editing, Conceptualization. **SH:** Writing - Review & Editing, Conceptualization. **AM:** Writing - Review & Editing, Conceptualization, Supervision. **AW:** Conceptualization, Supervision, Resources, Funding acquisition. **JBB:** Writing - Review & Editing, Conceptualization, Supervision, Resources, Funding acquisition. **BvdW:** Writing - Review & Editing, Conceptualization, Supervision, Resources, Funding acquisition.

## Notes

### Competing Interest Statement

The authors have declared no competing interest.

## References

Bischoff LJM, Kuijper IA, Schimming JP, Wolters L, Braak B ter, Langenberg JP, Noort D, Beltman JB, van de Water B. 2019. A systematic analysis of Nrf2 pathway activation dynamics during repeated xenobiotic exposure. Arch Toxicol. 93(2):435–451. doi:10.1007/S00204-018-2353-2.

ter Braak B, Klip JE, Wink S, Hiemstra S, Cooper SL, Middleton A, White A, van de Water B. 2022. Mapping the dynamics of Nrf2 antioxidant and NFκB inflammatory responses by soft electrophilic chemicals in human liver cells defines the transition from adaptive to adverse responses. Toxicology in Vitro. 84:105419. doi:10.1016/J.TIV.2022.105419.

Copple IM, den Hollander W, Callegaro G, Mutter FE, Maggs JL, Schofield AL, Rainbow L, Fang Y, Sutherland JJ, Ellis EC, et al. 2019. Characterisation of the NRF2 transcriptional network and its response to chemical insult in primary human hepatocytes: implications for prediction of drug-induced liver injury. Arch Toxicol. 93(2):385–399. doi:10.1007/S00204-018-2354-1.

Di Z, Herpers B, Fredriksson L, Yan K, van de Water B, Verbeek FJ, Meerman JHN. 2012. Automated Analysis of NF-κB Nuclear Translocation Kinetics in High-Throughput Screening. PLoS One. 7(12). doi:10.1371/journal.pone.0052337.

Hafner A, Bulyk ML, Jambhekar A, Lahav G. 2019. The multiple mechanisms that regulate p53 activity and cell fate. Nat Rev Mol Cell Biol. 20. doi:10.1038/s41580-019-0110-x.

Hanlon N, Coldham N, Gielbert A, Kuhnert N, Sauer MJ, King LJ, Ioannides C. 2008. Absolute bioavailability and dose-dependent pharmacokinetic behaviour of dietary doses of the chemopreventive isothiocyanate sulforaphane in rat. Br J Nutr. 99(3):559–564. doi:10.1017/S0007114507824093.

Hetz C, Papa FR. 2018. The Unfolded Protein Response and Cell Fate Control. Mol Cell. 69(2):169–181. doi:10.1016/j.molcel.2017.06.017.

Hiemstra S, Ramaiahgari SC, Wink S, Callegaro G, Coonen M, Meerman J, Jennen D, van den Nieuwendijk K, Dankers A, Snoeys J, et al. 2019. High-throughput confocal imaging of differentiated 3D liver-like spheroid cellular stress response reporters for identification of drug-induced liver injury liability. Arch Toxicol. 93(10):2895–2911. doi:10.1007/s00204-019-02552-0.

Hong DS, Kurzrock R, Supko JG, He X, Naing A, Wheler J, Lawrence D, Eder JP, Meyer CJ, Ferguson DA, et al. 2012. A phase I first-in-human trial of bardoxolone methyl in patients with advanced solid tumors and lymphomas. Clin Cancer Res. 18(12):3396–3406. doi:10.1158/1078-0432.CCR-11-2703.

Hu C, Eggler AL, Mesecar AD, Van Breemen RB. 2011. Modification of Keap1 cysteine residues by sulforaphane. Chem Res Toxicol. 24(4):515–521. doi:10.1021/TX100389R.

Kubo E, Chhunchha B, Singh P, Sasaki H, Singh DP. 2017. Sulforaphane reactivates cellular antioxidant defense by inducing Nrf2/ARE/Prdx6 activity during aging and oxidative stress. Scientific Reports 2017 7:1. 7(1):1–17. doi:10.1038/s41598-017-14520-8.

Leist M, Ghallab A, Graepel R, Marchan R, Hassan R, Bennekou SH, Limonciel A, Vinken M, Schildknecht S, Waldmann T, et al. 2017. Adverse outcome pathways: opportunities, limitations and open questions. Arch Toxicol. 91(11):3477–3505.

Liby K, Hock T, Yore MM, Suh N, Place AE, Risingsong R, Williams CR, Royce DB, Honda T, Honda Y, et al. 2005. The Synthetic Triterpenoids, CDDO and CDDO-Imidazolide, Are Potent Inducers of Heme Oxygenase-1 and Nrf2/ARE Signaling. Cancer Res. 65(11):4789– 4798. doi:10.1158/0008-5472.CAN-04-4539.

Liu A, Han N, Munoz-Muriedas J, Bender A. 2022. Deriving time-concordant event cascades from gene expression data: A case study for Drug-Induced Liver Injury (DILI). PLoS Comput Biol. 18(6):e1010148. doi:10.1371/JOURNAL.PCBI.1010148.

Liu J, Wu KC, Lu Y-F, Ekuase E, Klaassen CD. 2013. NRF2 Protection against Liver Injury Produced by Various Hepatotoxicants. Oxid Med Cell Longev. 2013(305861). doi:10.1155/2013/305861.

Mathew ST, Bergström P, Hammarsten O. 2014. Repeated Nrf2 stimulation using sulforaphane protects fibroblasts from ionizing radiation. Toxicol Appl Pharmacol. 276(3):188–194. doi:10.1016/j.taap.2014.02.013.

Morgenstern C, Lastres-Becker I, Demirdöğen BC, Costa VM, Daiber A, Foresti R, Motterlini R, Kalyoncu S, Arioz BI, Genc S, et al. 2024. Biomarkers of NRF2 signalling: Current status and future challenges. Redox Biol. 72:103134. doi:10.1016/J.REDOX.2024.103134.

Poser I, Sarov M, Hutchins JRA, Hériché JK, Toyoda Y, Pozniakovsky A, Weigl D, Nitzsche A, Hegemann B, Bird AW, et al. 2008. BAC TransgeneOmics: A high-throughput method for exploration of protein function in mammals. Nat Methods. 5(5):409–415. doi:10.1038/nmeth.1199.

Sharma RP, Loonstra-Wolters L, ter Braak B, Niemeijer M, White A, van de Water B, Middleton AM, Beltman JB. 2025. Deciphering the quantitative relationship between NRF2 and SRXN1 through semi-mechanistic computational modeling. Submitted.

Ramaiahgari SC, Den Braver MW, Herpers B, Terpstra V, Commandeur JNM, Van De Water B, Price LS. 2014. A 3D in vitro model of differentiated HepG2 cell spheroids with improved liver-like properties for repeated dose high-throughput toxicity studies. Arch Toxicol. 88(5):1083–1095. doi:10.1007/s00204-014-1215-9.

Souza TM, Kleinjans JCS, Jennen DGJ. 2017. Dose and time dependencies in stress pathway responses during chemical exposure: Novel insights from gene regulatory networks. Front Genet. 8(OCT):288554. doi:10.3389/FGENE.2017.00142/BIBTEX.

Takaya K, Suzuki T, Motohashi H, Onodera K, Satomi S, Kensler TW, Yamamoto M. 2012. Validation of the Multiple Sensor Mechanism of the Keap1-Nrf2 System. Free Radic Biol Med. 53(4):817. doi:10.1016/J.FREERADBIOMED.2012.06.023. [accessed 2025 Jun 23]. https://pmc.ncbi.nlm.nih.gov/articles/PMC3539416/.

Thimmulappa RK, Mai KH, Srisuma S, Kensler TW, Yamamoto M, Biswal S. 2002. Identification of Nrf2-regulated Genes Induced by the Chemopreventive Agent Sulforaphane by Oligonucleotide Microarray. Cancer Res. 62:5196–5203.

Vinken M. 2013. The adverse outcome pathway concept: A pragmatic tool in toxicology. Toxicology. 312(1):158–165. doi:10.1016/J.TOX.2013.08.011.

Weaver RJ, Blomme EA, Chadwick AE, Copple IM, Gerets HHJ, Goldring CE, Guillouzo A, Hewitt PG, Ingelman-Sundberg M, Jensen KG, et al. 2020. Managing the challenge of drug-induced liver injury: a roadmap for the development and deployment of preclinical predictive models. Nat Rev Drug Discov. 19. doi:10.1038/s41573-019-0048-x.

Wei J, Zhao Q, Zhang Y, Shi W, Wang H, Zheng Z, Meng L, Xin Y, Jiang X. 2021. Sulforaphane-Mediated Nrf2 Activation Prevents Radiation-Induced Skin Injury through Inhibiting the Oxidative-Stress-Activated DNA Damage and NLRP3 Inflammasome. Antioxidants. 10(11). doi:10.3390/ANTIOX10111850.

Wijaya LS, Gabor A, Pot IE, van de Have L, Saez-Rodriguez J, Stevens JL, L. Dévédec SE, Callegaro G, Van de Water B. 2024. A network-based transcriptomic landscape of HepG2 cells uncovering causal gene-cytotoxicity interactions underlying drug-induced liver injury. Toxicol Sci. 198(1):14–30. doi:10.1093/TOXSCI/KFAD121.

Wink S, Burger GA, L. Dévédec SE, Beltman JB, Van De Water B. 2022. User-friendly high-content imaging analysis on a single desktop: R package H5CellProfiler. BioRxiv. doi:10.1101/2022.10.06.511212.

Wink S, Hiemstra S, Herpers B, van de Water B. 2017. High-content imaging-based BAC-GFP toxicity pathway reporters to assess chemical adversity liabilities. Arch Toxicol. 91(3):1367–1383.

Wink S, Hiemstra S, Huppelschoten S, Danen E, Niemeijer M, Hendriks G, Vrieling H, Herpers B, van de Water B. 2014. Quantitative high content imaging of cellular adaptive stress response pathways in toxicity for chemical safety assessment. Chem Res Toxicol. 27(3):338–355.

Wink S, Hiemstra SW, Huppelschoten S, Klip JE, van de Water B. 2018. Dynamic imaging of adaptive stress response pathway activation for prediction of drug induced liver injury. Arch Toxicol. 92(5):1797–1814.

Yamamoto M, Kensler TW, Motohashi H. 2018. The KEAP1-NRF2 system: A thiol-based sensor-effector apparatus for maintaining redox homeostasis. Physiol Rev. 98(3):1169–1203. doi:10.1152/PHYSREV.00023.2017.

